# The engagement of layer 6 corticothalamic neurons in somatosensation

**DOI:** 10.64898/2026.08.11.744301

**Authors:** Elaida D Dimwamwa, Nelson H Chang, Christian Waiblinger, Garrett B Stanley

**Author notes:** **Corresponding author:** Professor Garrett B Stanley, Coulter Department of Biomedical Engineering, Georgia Institute of Technology & Emory University, 313 Ferst Drive, Atlanta GA 30332-0535 USA.

## Abstract

The corticothalamic neurons from layer 6 (L6CT) of primary sensory cortices provide extensive input to the thalamus in addition to projecting across layers intracortically, positioning them to play a key hypothesized role in shaping thalamocortical signaling. With the expansion of tools for precise functional identification of L6CT neurons during in-vivo electrophysiology, increasing evidence highlights L6CT neurons as dynamic gain modulators of thalamocortical sensory responses. However, much of the work to date has been conducted under anesthesia and not in the context of awake and/or behaving animals. In this study, we show in the awake mouse that L6CT neurons convey information about the ascending sensory input, the timing of which is fast enough to contribute to the sensory response of neurons throughout the thalamocortical circuit. Overall, L6CT neurons robustly encode the presence vs absence of a sensory stimulus but are relatively weak encoders of the fine details. Benchmarked against the activity of other excitatory cortical neurons, we also provide evidence for L6CT neurons as predictors of behavioral outcome during a trained detection task. Taken together, the results in this study potentially tie L6CT neurons to behavior in tactile detection, one of the most fundamental functional roles of the pathway.

**Significance Statement:** Layer 6 corticothalamic (L6CT) neurons provide one of the major feedback pathways from the sensory cortex to the thalamus, yet the extent to which their role in sensory processing remains poorly understood. We show that L6CT neurons in mouse somatosensory cortex robustly signal the presence of tactile stimuli but are comparatively weak encoders of stimulus strength. During a sensory detection task, their responses strongly reflect whether the animal detects the stimulus and predict behavioral outcome better than other cortical excitatory neurons. These findings suggest that L6CT neurons are specialized less for representing fine sensory detail and more for signaling the occurrence and behavioral relevance of sensory events, providing insight into when and how corticothalamic feedback participates in perception.

## Introduction

Motivated by the striking density of the projections of cortical layer 6 corticothalamic (L6CT) neurons onto thalamic neurons, in 1906, Ramón y Cajal asked whether L6CT neurons are “indispensable” for the propagation of sensory information through the processing hierarchy (Ramón Y Cajal, 1906). Since then, a plethora of investigations across sensory systems have dissected the complex nature of L6CT neuronal inputs on their thalamic and cortical targets (Augustinaite & Kuhn, 2020; Crandall et al., 2015; McCormick & von Krosigk, 1992; Sherman & Koch, 1986; Van Horn et al., 2000; von Krosigk et al., 1999) and have established L6CT neurons as excitatory “modulators,” not “drivers,” of thalamic neurons (Kirchgessner et al., 2021; Sherman & Guillery, 1998). Undoubtedly, the complexity of L6CT function is shaped by its anatomical projections: L6CT neurons provide monosynaptic excitatory inputs to both the other cortical neurons and thalamic neurons which in turn projects to the cortex; they also provide disynaptic inhibitory inputs to the thalamus via axon collaterals to the inhibitory thalamic reticular nucleus which in turn projects to the thalamus. Various studies have uncovered the myriad of ways in which L6CT neurons can dynamically shape activity in thalamocortical circuits. This ranges from L6CT-mediated shifts in the gain of sensory inputs (Bortone et al., 2014; Crandall et al., 2015; Cudeiro et al., 2000; Denman & Contreras, 2015; Dimwamwa et al., 2024; W. Guo et al., 2017; Kirchgessner et al., 2020, 2021; Olsen et al., 2012; Reinhold et al., 2023; Sillito et al., 1994; Voigts et al., 2020; Ziegler et al., 2023), the reliability of sensory responses (Mease et al., 2014; Pauzin & Krieger, 2018), the timing of sensory responses (Hasse & Briggs, 2017), spatial integration (Andolina et al., 2007; Born et al., 2021; Eyding et al., 2003; Temereanca et al., 2008; W. Wang et al., 2016), and selective encoding of deviant sensory inputs (Voigts et al., 2020). However, this evidence must contend with the many observations demonstrating that L6CT neurons are incredibly sparsely firing neurons. Despite the location of L6CT neurons in primary sensory cortices, reports of sensory responses much less than 1 spikes/sec are numerous, as are reports of very low spontaneous firing rates (Kwegyir-Afful & Simons, 2009; Lee et al., 2008; Swadlow, 1987, 1989; Swadlow & Hicks, 1996; Tsumoto & Suda, 1980; Vélez-Fort et al., 2018). While it is well established that L6CT neurons receive both bottom-up sensory inputs as well as top-down inputs from a variety of sensory and extra-sensory regions (W. Guo et al., 2017; Whilden et al., 2021), these findings raise questions about the contexts in which L6CT neurons *would* modulate sensory processing.

Importantly, of the few direct measurements of L6CT activity, the majority were conducted either in brain slices or in an anesthetized *in vivo* preparation, both of which would impact the ability to measure ongoing and stimulus-evoked spiking in physiological and behaviorally relevant conditions. L6CT neurons receive direct thalamic inputs (Constantinople & Bruno, 2013; Deschênes et al., 1998) so one may expect that they would encode bottom-up inputs for transmission throughout the thalamocortical circuit. Yet, the motor and cholinergic inputs to L6CT neurons suggests an extra-sensory contribution of L6CT neurons to sensory processing. However, the exact relation of L6CT neuron activity to basic sensory processing and perception remains enigmatic.

To better understand of the role of L6CT neurons in sensation, we targeted silicon probe recordings to the primary somatosensory cortex (S1) in awake mice and compared the activity of optogenetically identified L6CT neurons to that of other excitatory S1 neurons. We find that L6CT neurons are indeed sensory-responsive in the awake animal and that their activity reliably encodes the presence, more so than the magnitude, of a sensory stimulus. Motivated by this finding, we then measured L6CT neuron activity in mice behaving in a sensory detection task and found that L6CT neuron sensory responses encode the behavioral outcome of the animal better than other S1 neurons. Collectively, these data provide evidence linking L6CT neuron activity to its previously hypothesized role in the thalamocortical gating of sensory inputs.

## Materials and Methods

### Experimental model and subject details

Experimental subjects consisted of five female and six male NTSR1-cre mice (B6.FVB(Cg)-Tg(Ntsr1-cre)GN220Gsat/Mmcd, MMRRC; Gong et al., 2003, 2007) aged between seven and 19 weeks at the start of experimentation. Mice were housed under a reversed light-dark cycle and, when possible, in sibling pairs. All procedures were approved by the Institutional Animal Care and Use Committee at the Georgia Institute of Technology and adhered to guidelines established by the National Institutes of Health.

### Animal preparation

The mice were first implanted with a custom, stainless steel headplate while anesthetized with isoflurane anesthesia (1-1.5%). The mice were given a minimum of three days to recover (Bolus et al., 2018; Borden et al., 2022; Dimwamwa et al., 2024; Z. V. Guo et al., 2014; Liew et al., 2021, 2025; Pala & Stanley, 2022; Wright et al., 2021). Then, an image of the superficial vasculature overlaid with a map of the whisker columns of the primary somatosensory cortex (S1) was generated via intrinsic signal optical imaging conducted under 1-1.5% isoflurane anesthesia through a thinned skull (Dimwamwa et al., 2024; Masino et al., 1993; Pala & Stanley, 2022).

### AAV injection

At least three weeks prior to the electrophysiological recordings, the mice were injected with 800 nL of AAV5-EF1a-DIO-hChR2(H134R)-eYFP.WPRE.hGH (UPenn Vector Core). The virus was injected into the C1 whisker column based on the map generated via intrinsic signal optical imaging. The injection depth was 800 um. The mice were then provided a minimum of three days to recover (Dimwamwa et al., 2024).

### Whisker stimulation

Sensory inputs consisted of the precise deflection of an individual whisker contralateral to the electrophysiological recording site. The whisker was threaded into a narrow, 1.5 cm long tube and deflected in the rostral-caudal plane using a galvo-motor (galvanometer optical scanner model 6210H, Cambridge Technology). The galvanometer was controlled at 1 kHz by custom scripts written in MATLAB and Simulink Real-Time (MathWorks, Natick, MA; Borden et al., 2022; Dimwamwa et al., 2024; Liew et al., 2025; Pala & Stanley, 2022; Wright et al., 2021). The whiskers were deflected using a 10 ms long cosine waveform of varying strengths. The waveform strength is defined by the amplitude – 0, 2, 4, 8, and 16 deg – which correspond to mean velocities – 0, 204, 408, 816, and 1631 deg/s - to which thalamocortical neurons are most sensitive (Waiblinger et al., 2015, 2022).

### Behavioral paradigm and training

The mice were trained in a standard Go-No-Go detection task (Ollerenshaw et al., 2012; Waiblinger et al., 2019). For this task, water-restricted and head-restrained mice ultimately learned to lick for a water reward upon deflection of individual whiskers. The training process for each mouse was as follows. After a few days of habituating the mouse to the recording setup and the lick spout, the initial phase of learning would begin, the goal being for the mouse to learn the association between the stimulus and the reward. In this phase, trials consisted of water reward being automatically given with each whisker deflection. The whisker deflection strength was 16 deg, the largest of all strengths ultimately considered. 80% of trials had a stimulus, while the remaining 20% were “catch” trials, or trials in which there is no sensory input (0 deg). The inter-trial interval was between 6 and 8 seconds, drawn randomly from a uniform distribution for each trial. Regardless of the training phase, sessions were stopped by the experimenter once the animal did not lick in 10 consecutive trials. The body weight was monitored daily and supplemental water was provided as necessary. For two days of the week, behavioral training was paused and the mouse was given unlimited access to water in its home cage.

Once the animal consistently licked in stimulus trials (typically within less than five sessions), the mouse was transitioned to the next phase of learning. In this phase, to be rewarded, the mouse was required to first lick within one second of each stimulus presentation, hereafter referred to as the response window. In addition, there was a “no-lick” window that lasted for the second before each stimulus presentation. If the mouse licked during the no-lick window, the trial was aborted. But if the animal did not lick in the no-lick window, the trial would continue and result in one of four possible outcomes based on the mouse’s behavior. In the case where a stimulus was presented, a trial is defined as a “hit” if the mouse successfully licked for reward within the response window and a “miss” if the animal did not. In the case of a catch trial, the trial was defined as a “false alarm” if the animal licked in the response window and a “correct rejection” if the animal did not.

For each session, the learning of the mouse was assessed by computing *d’*^′^, or the effect size from the observed hit rate and false alarm rate:

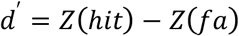

where *Z*(ℎ*it*) and *Z*(*fa*) are the inverse cumulative distribution function of the gaussian distribution of the hit and false alarm rates, respectively. Animals were considered expert when they performed at a minimum *d*^′^of 2.3, calculated with a hit rate of 0.95 and false alarm rate of 0.25 (Carandini & Churchland, 2013; Waiblinger et al., 2022). This typically occurred within 3-5 weeks.

Once having reached expert level, stimuli of one of five strengths were presented - 0, 2, 4, 8, and 16 degrees - throughout the session. The stimuli were presented in a pseudorandom order such that within each block of five trials, the order in which each stimulus strength was presented was randomized; the stimuli were drawn from a uniform distribution.

During learning, the task began as soon as the mouse was head-restrained. But once the mouse reached expert level, the animal was habituated to waiting while head-restrained, before the task would begin. Over the course of four to five sessions, the wait time was increased from five minutes to one hour. This wait time was to prepare the mouse for the electrophysiological recordings, where we first inserted the probe and waited for the brain to settle before beginning electrophysiological recording and allowing the mouse to begin the task. During these habituation sessions, the mouse was also habituated to flashes of an LED, necessary for optogenetic stimulation (see below).

All behavioral sessions were conducted in darkness with ambient white noise to mask the potential sound of the whisker galvanometer. To confirm that the mouse was using somatosensory input and not visual or auditory cues to perform the task, control sessions were conducted. In these sessions, after the mouse completed 50 trials in the task, 50 additional trials were conducted where the whisker was removed from the galvanometer. The galvanometer was placed near the whisker pad, but not connected to an individual whisker. We verified that the behavioral performance of all animals dropped to chance once the whisker was removed. The animals could otherwise perform the task regardless of which whisker was being deflected.

### Electrophysiological recordings

All electrophysiological recordings were performed in head-restrained mice either under isoflurane anesthesia (1-1.5%) or during wakefulness. In either case, a small craniotomy to target an S1 whisker column contralateral to the side of the face provided with sensory input was made while the mouse was under isoflurane anesthesia (1-1.5%). The targeted location of the craniotomy was based on the previously generated intrinsic optical signal map (see “Animal preparation”). For anesthetized recordings, we then immediately proceeded to insert the recording probe. The animal was maintained at 1-1.5% isoflurane for the remainder of the recording. For awake recordings, the craniotomy was covered with silicone elastomer (Kwik-Cast, WPI, Sarasota, FL) and the mice were given no less than 2 hours to recover before being head-restrained, the elastomer removed, and the recording probes inserted.

Both the anesthetized and awake electrophysiological recordings were performed with 64 channel silicon probes (A1x64-Poly2-6mm-23s-160 or A1x64-Edge-6mm-20-177-A64, both from Neuronexus, Ann Arbor, MI). For one session per craniotomy, the probe was coated with DiI (0.2 mg/mL in ethanol, Invitrogen, Waltham, MA) to enable fluorescence imaging of the probe track (see “Histological verification” below). The probes were then inserted at a 35 degree angle to depths between 1250 and 1400 μm over a period of approximately 10 minutes (Dimwamwa et al., 2024; Pala & Stanley, 2022). The brain tissue was given approximately 30 minutes to settle before data acquisition began. During this waiting period, we manually deflected the whiskers and determined which whisker elicited the largest neuronal response in the column in which our probe was inserted. This whisker is referred to as the primary whisker. The primary whisker was determined for each recording session and was used to deliver sensory input for that session.

The electrophysiological signals were acquired using the Cerebus acquisition system (Blackrock Neurotech, Salt Lake City, Utah). The signals were amplified, filtered between 0.3 Hz and 7.5 kHz, and digitized at 30,000 Hz.

One recording was performed per day, no more than two recordings were conducted per craniotomy, and up to three craniotomies were made per mouse.

### Optogenetic stimulation

To identify L6CT neurons, 470 nm light emitting diode (LED) outputs were delivered via a 400 μm diameter optical fiber placed near the recording probe, on the skull surface. The LED output was controlled at a resolution of 1 kHz with custom scripts written in MATLAB and Simulink Real-Time (MathWorks, Natick, MA). LED waveforms consisted of rectangular pulses that were of 30 mW/mm^2^ intensity and 5 ms in duration. The interval between LED pulses was between 0.5 and 1 sec, drawn from a uniform distribution.

### Whisker videography and whisker movement analysis

During the electrophysiological recording sessions conducted in awake mice, the movements of whiskers on both sides of the face were captured using a CCD camera (EoSens CL MC1362, Mikrotron, Germany) that acquired frames at 200 Hz. Facemap (Syeda et al., 2022) was used to extract motion energy from manually defined regions of interest (ROIs) around both sides of the face. Custom scripts were then written to determine whether the whiskers were moving in each frame. Briefly, a time series of the recording session was created by taking the summed squared motion energy across all ROIs in each frame. Time points in this time series that were above 3.5 times the standard deviation of the entire time series were labelled to belong to epochs of whisker movement. In all analyses of neuronal sensory responses, only trials with no whisker movement 75 ms before and after each sensory stimulus were included. In all analyses of ongoing neuronal activity, only trials with no whisker movement during the window of interest were included (Dimwamwa et al., 2024; Pala & Stanley, 2022).

### Histological verification

Expression of ChR2 in L6CT neurons as well as the location of the recording probe track coated in DiI (0.2 mg/mL in ethanol, Invitrogen, Waltham, MA) was verified with fluorescence imaging (DiCarlo et al., 1996; Dimwamwa et al., 2024; Liew et al., 2025; Pala & Stanley, 2022). Briefly, after the final behavioral session, the mice were transcardially perfused with 1x PBS (137 mM NaCl, 2.7 mM KCl, and 10 mM PB, VWR), followed by 4% PFA. The brains were extracted, post-fixed in 4% PFA, and then sectioned to 100 um-thick coronal slices using a vibratome (VT1000S, Leica Biosystems, Deer Park, IL). The slices were then incubated with DAPI (2 μm in PBS) for 15 min, rinsed in PBS, mounted on slides with DABCO solution (1,4-Diazabicyclo[2.2.2]octane, Sigma), and imaged under a confocal microscope (Laser Scanning Microscope 900, Zeiss, Germany).

### Spike-sorting and S1 neuron classification

The 64 channel electrophysiological signals were high-pass filtered (3^rd^ order Butterworth filter with a 500 Hz cutoff frequency) and median-filtered across channels before being processed through Kilosort for template matching-based, automatic spike sorting (Pachitariu et al., 2024). The extracted clusters were then manually curated using Phy2 (Rossant & Harris, 2013) to remove non-neuronal clusters. Cluster quality metrics were then extracted for the remaining clusters using custom scripts in MATLAB. All neuron clusters considered for analysis had at least 300 spikes, a signal-to-noise ratio of the mean spike waveform greater than three, a mean waveform voltage from peak to trough greater than 40 μV, a coefficient of variation of the spiking over the duration of the recording less than 1 (using 120-second-long bins), and less than 2% of all spikes violating the 2 ms refractory period. Note that the nature of any claims presented does not change with more strict cluster quality criteria.

### Cortical cell type classification

S1 neurons were classified as regular spiking (RS) or fast spiking (FS) based on the width of their mean spike waveform, defined as the trough to peak time (TPT). Neurons with a TPT greater than 0.5 ms were classified as RS neurons and those with a TPT less than 0.4 were classified as FS neurons. Neurons with TPT’s in between 0.4 and 0.5 were omitted from analysis (Dimwamwa et al., 2024; Pala & Stanley, 2022).

### Cortical layer assignment

Each neuron was assigned to a cortical layer based on the channel on which its mean waveform from trough-to-peak was largest. Layer assignment of each channel was based on a functional estimate of the center of layer 4. Specifically, we extracted the trial-averaged sensory response for each channel on the recording probe with the following data types: threshold-crossings of the high-pass filtered recording trace, the local field potential (LFP), and current-source density (CSD) analysis. The center of layer 4 was determined based on collective evidence of the channels that had the earliest onset and largest responses across these data types as well as a sink in the CSD analysis. All channels were then assigned to a layer based on their distance from the assigned center of layer 4 (Dimwamwa et al., 2024; Pala & Stanley, 2022; Sederberg et al., 2019; Sofroniew et al., 2015) (Sofroniew et al., 2015; Pala and Stanley, 2022; Dimwamwa et al., 2024). Each layer was assigned the appropriate number of channels according to the following thicknesses: layer 2/3: 300 μm; layer 4: 200 μm; layer 5: 300 μm; layer 6: 300 μm (Dimwamwa et al., 2024; Hooks et al., 2011; Pala & Stanley, 2022; Sederberg et al., 2019).

### Identification of L6CT neurons

L6CT neurons were identified based on quantifications of their across-trial response to 5 ms long optogenetic LED pulses. Only deep, RS S1 neurons that were LED-responsive (see below) were considered as candidate L6CT neurons. Given expression of ChR2 specific to the L6CT neurons (Gong et al., 2007; W. Guo et al., 2017), identification of L6CT neurons from blind, extracellular electrode recordings was based on whether the LED pulse causes a significant increase in each neuron’s spiking compared to spontaneous spiking, with little temporal jitter across trials. This assessment was performed using the stimulus-associated spike latency test (SALT; Kvitsiani et al., 2013). Using this approach, any deep RS S1 neuron with a p-value less than 0.01 was classified as a L6CT neuron.

### Analysis of neuronal activity

#### Classifying neurons as LED-responsive

Neurons were classified as LED-responsive based on the two-sided Wilcoxon signed rank test applied to the trial-by-trial spike counts in a 20 ms window before the 5 ms LED pulse compared to spike counts in an equivalent window after the LED pulse.

#### Classifying neurons as sensory-responsive

To be classified as sensory-responsive, neurons were required to pass a minimum of two out of three tests assessing whether the stimulus resulted in a statistically significant increase in spiking from baseline spiking. The first test was significance as determined by the two-sided Wilcoxon signed rank test on the trial-by-trial spike counts in a 50 ms window before the sensory stimulus compared to spike counts in an equivalent window after the sensory stimulus. The second test assessed whether zero overlapped with the 95% bootstrapped confidence interval generated from the baseline subtracted spike counts in the 50 ms post-stimulus window. The third test assessed whether a minimum of three bins from a 10 ms resolution PSTH were above or two bins of the PSTH were below the 95% bootstrapped confidence interval of pre-stimulus activity (Dimwamwa et al., 2024; Pala & Stanley, 2022).

#### Measuring sensory responses

Sensory responses were quantified by calculating the average rate of spikes across trials in a 30 ms post-stimulus window. These rates were subtracted on a trial-by-trial basis by the rate of spontaneous spikes in the 30 ms pre-stimulus window. To determine whether a neuron was sensory-responsive, the trial-by-trial pre- and post-stimulus spike rates in the same 30 ms windows were compared using the two-sided Wilcoxon signed rank test. Neurons with a significant change in post-stimulus spike rates were considered sensory-responsive.

#### Measuring response latencies

To measure the response latency to either sensory or LED inputs, we assessed the first spike latency, or the average time post-stimulus of the first spike in trials. For the sensory response latency, the spike must occur within 30 ms of the sensory input and for the LED response latency, the spike must occur within 20 ms of the LED input.

### Linear discriminant analysis (LDA) decoding and spike count parametric fits

To determine the extent to which various features (e.g. the sensory stimulus strength or the animal’s behavioral outcome) could be decoded from a population’s neuronal activity, we performed LDA of the population’s activity. Because both the number of simultaneously recorded L6CT neurons was limited and also the number of recorded other RS neurons far outweighs the number of recorded L6CT neurons, it was necessary to perform this estimation using all recorded neurons of each particular subtype, across all recording sessions and mice. We randomly drew with replacement the spiking from 25 trials for 25 neurons from the neuron population being analyzed. The joint spike counts from 80% of the trials for the subset of neurons was used to train an LDA classifier using MATLAB’s *fitcdiscr* function. The classifier’s performance was assessed on the remaining 20% of trials using the *predict* function. This process was iterated 1000 times to comprehensively assess the information contained within the respective neuron population ((Liew et al., 2025; Pala & Stanley, 2022; Q. Wang et al., 2010). Varying the number of sampled neurons and trials did not affect the nature of the results. Additionally, use of a Bayesian decoder produced similar results.

For visualization purposes, probability density functions (pdfs) were generated for each iteration by fitting a gamma distribution to the joint spike counts of the subset of neurons in each trial. The mean, *μ*, and the variance, *σ*^2^, were extracted from the joint spike counts across trial. Then, the two parameters defining a gamma distribution, *α*, the shape parameter, and *Θ*, the scale parameter, were computed as follows:

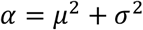

and

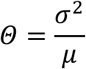

The *gampdf* function in MATLAB was then used to generate the pdfs.

### Statistical analyses

To test for significant changes in measures of neuronal activity, the two-sided Wilcoxon signed rank test with a Bonferroni correction was used between conditions/populations. To determine whether there was a monotonic trend in the data, the Spearman’s rank correlation test was used. All tests were implemented using built-in MATLAB functions.

### Data availability

The processed data generated in this study will be made publicly available upon acceptance of the research article, via Zenodo, and linked here.

## Results

To characterize the activity of L6CT neurons in the awake brain, we performed silicon probe recordings of the spiking activity of populations of individual neurons throughout the depth of the primary somatosensory cortex (S1) in awake and head-fixed NTSR1-cre mice. The NTSR1-cre line selectively expresses cre-recombinase in corticothalamic neurons of neocortical layer 6 (Gong et al., 2003, 2007). The mice were injected with a recombinant adeno-associated virus encoding cre-dependent channelrhodopsin (ChR2) fused with eYFP, enabling us to functionally identify L6CT neurons from extracellular recordings. The selective ChR2 expression in the processes of L6CT neurons was verified with confocal fluorescence microscopy (Figure 1a, see Methods). To identify L6CT neurons from our recordings, we presented 5 ms, 470 nm LED pulses via an optical fiber positioned at the S1 cortical surface. Current source density (CSD) analysis of the average response to the 5 ms LED pulse reveals an early sink in layer 6, indicative of activation of the neurons in layer 6 (Figure 1b). L6CT neurons were ultimately optotagged amongst all putative excitatory, regular-spiking (RS) neurons on the basis of their spiking response to the LED pulse (see Methods). Specifically, RS neurons were identified as neurons with waveform width from trough to peak greater than 0.4 msec (Figure 1c; Barthó et al., 2004). L6CT neurons were then differentiated from other RS neurons using the stimulus-associated spike latency test (SALT), a statistical method to determine whether the LED pulse drove a rapid and significant increase in the spiking of individual neurons with low temporal jitter compared to spontaneous spiking (see Methods; Kvitsiani et al., 2013). Importantly, the SALT-positive, putative L6CT neurons originated either within layer 6 or within a few channels of the functionally identified layer border (n = 44 SALT-positive L6CT neurons). The SALT-negative but LED-responsive RS neurons, which are presumably polysynaptically connected to L6CT neurons, were recorded throughout the cortical depth (n = 80 SALT-negative RS neurons). Figure 1d shows the grand-average peri-stimulus time histogram (PSTH) of the recorded population of optotagged putative L6CT neurons and all other recorded SALT-negative RS neurons whose spiking was significantly enhanced by the LED pulse. The PSTH makes clear that beyond not being as well driven by the LED, the other SALT-negative RS neurons respond at longer latencies than the L6CT neurons. To quantify this, we measured the average first spike latency as well as the first spike jitter, or the standard deviation of the first spike. Compared to other RS neurons that are LED-responsive, L6CT neurons have short latency responses and the responses have low temporal jitter (Figures 1e & 1f; <u>Dimwamwa et al., 2024; Williamson & Polley, 2019)</u>. Note that optogenetic stimulation was used only for identification of L6CT neurons; the remainder of the data presented here represent naturally occurring, and not artificially driven, neuronal activity.

**Figure 1.**
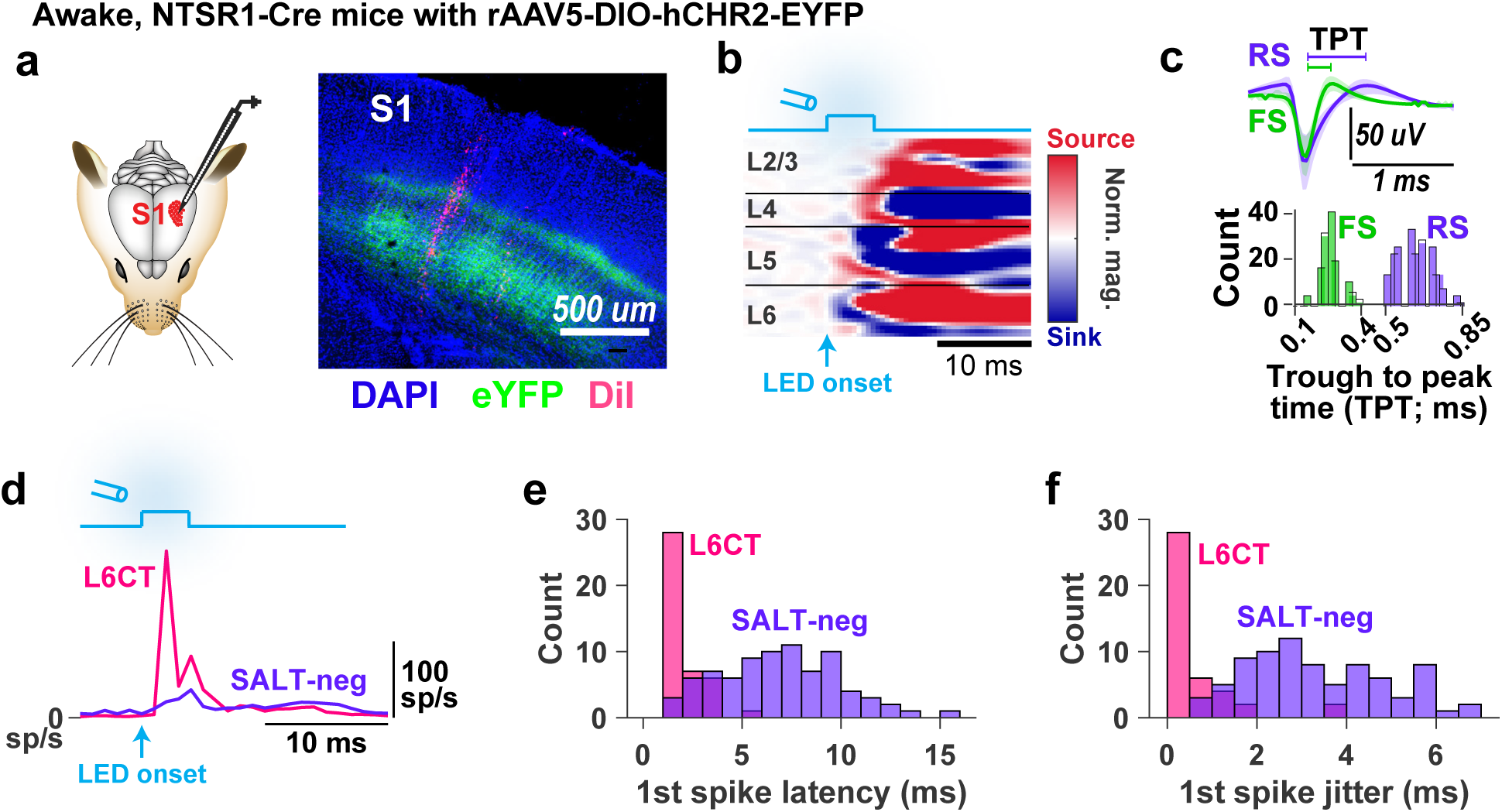
Optogenetic identification of L6CT neuron activity *in vivo*. **a.** Neuronal activity throughout the entire cortical depth of S1 was recorded using silicon probes in NTSR1-cre mice. The NTSR1-Cre mice were injected with a virus to selectively express ChR2-eYFP in L6CT neuron processes, as verified by coronal sections of DAPI labelled neurons (blue) and selective ChR2-eYFP expression in L6CT neuron processes (green). The recording probe track is labelled with DiI (magenta). **b.** Current source density (CSD) analysis of the cortical response in an example recording to a 5 ms LED pulse presented to an optical fiber at the cortical surface demonstrates the sink in L6. **c**.Top: the mean +/- standard deviation of all record ed neurons that are identified as either regular spiking (RS) or fast spiking (FS) based on the trough-to-peak time (TPT) of their mean waveform (n = 218 RS neurons and 159 FS neurons across 7 mice, 33 recordings). Bottom: the TPT of RS neurons is greater than 0.5 ms and that of FS neurons is less than 0.4 ms (see Methods). **d.** Grand PSTHs of putative L6CT neurons (n = 44 neurons, SALT-positive) and all other cortical SALT-negative RS neurons whose spiking is significantly enhanced by the LED pulse (excluding L6CT neurons; n = 80 neurons). **e.** Distribution of the average first spike latency for each neuron population presented in ***d*** in response to the LED pulse (see Methods). **f.** Same as ***e***, but for the first spike jitter, or the standard deviation of the first spike across successful trials (see Methods).

### L6CT neurons are sensory-responsive in awake animals

Having identified putative L6CT neurons *in vivo*, we then characterized the spiking activity of this cortical sub-population. Given the numerous previous reports of sparse or entirely absent sensory responses in L6CT neurons (Augustinaite & Kuhn, 2020; Kwegyir-Afful & Simons, 2009; Lee et al., 2008; Swadlow, 1987, 1989; Swadlow & Hicks, 1996; Tsumoto & Suda, 1980; Vélez-Fort et al., 2018) despite receiving direct inputs from the first-order thalamus, we sought to characterize the extent to which, if at all, S1 L6CT neurons are sensory-responsive during wakefulness. We recorded the spiking response of S1 neurons, including optotagged L6CT neurons, to brief, 16 deg deflections in the rostral-caudal plane of the individual whisker eliciting the largest and earliest response in the recorded cortical column (see Methods for details of whisker stimulation). We then quantified the baseline-subtracted sensory response. Given the known association of whisker movement to changes in thalamocortical spiking (Urbain et al., 2015), we exclusively analyzed trials in which there was no self-generated whisker movement as determined from analysis of videography of the mice’s whiskers (Figure 2a; see Methods). Figure 2b shows the sensory response of an example L6CT neuron during wakefulness; this neuron represents the 75^th^ percentile of all recorded L6CT neurons in terms of the magnitude of its sensory response. Notably, this neuron exhibits a relatively low spontaneous firing rate but is clearly sensory-responsive; this increase in spiking is at a relatively short latency following the stimulus, suggesting that the activity is induced from direct thalamic input. The sensory response of all recorded putative L6CT neurons is presented in Figure 2c, ordered from largest to smallest sensory-evoked response. Overall, we find that 53% (23 out of 44) of L6CT neurons have a statistically significant change in spiking from baseline and are thus sensory-responsive, with all responsive neurons increasing their spiking to the sensory stimulus and no neurons decreasing their spiking. Additionally, we measured an average response latency of 11.1 +/- 3.4 ms across L6CT neurons, which is slightly longer than layer 4 RS neurons whose average latency is 9.2 +/-1.4 ms.

**Figure 2.**
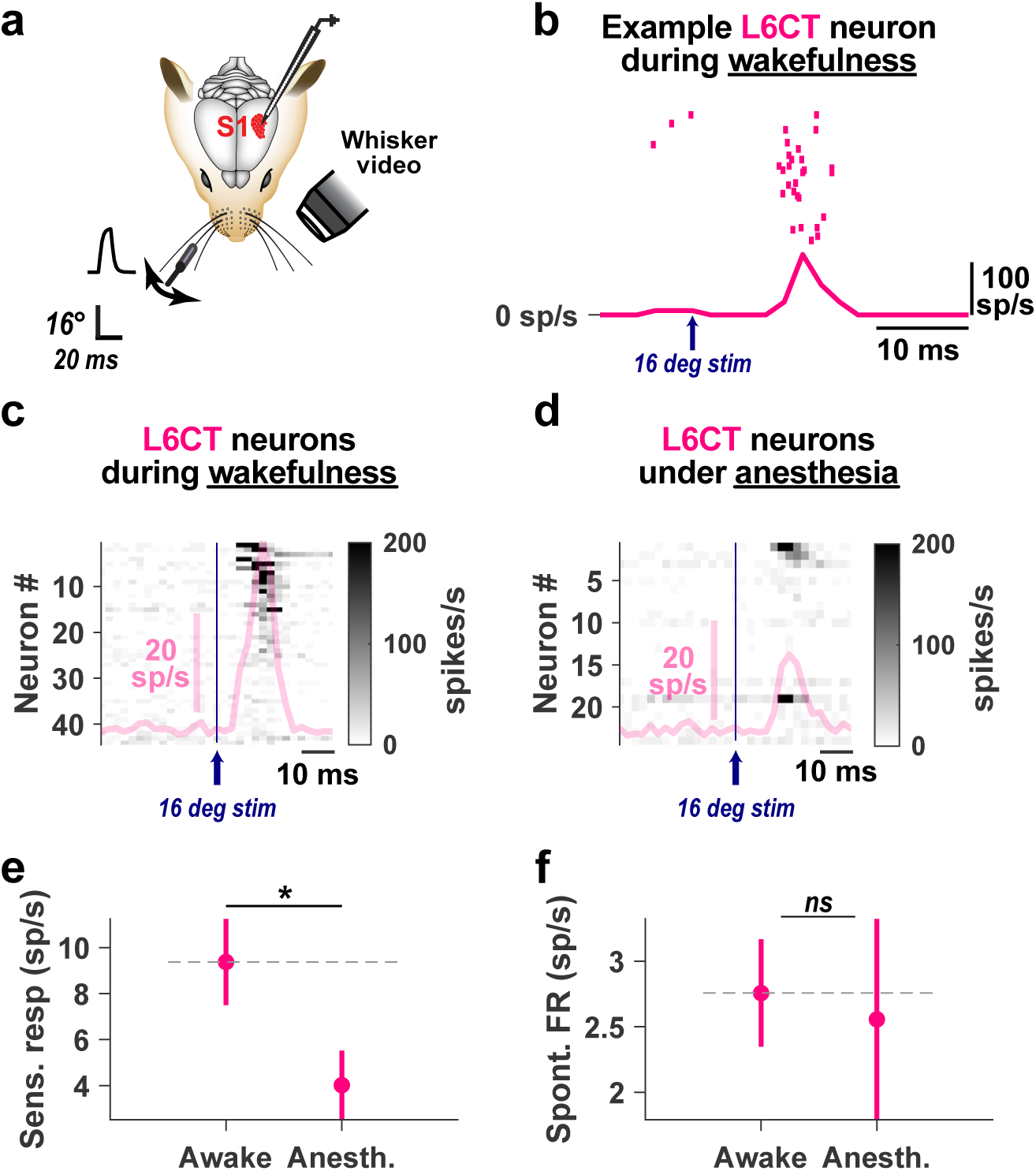
L6CT neurons are sensory-responsive in awake mice. **a.** Neuronal activity throughout the S1 cortical depth was recorded in NTSR1-cre mice expressing ChR2-eYFP while we deflected the individual whisker corresponding to the barrel column being recorded. The movement movement of the mouse’s whiskers was monitored via videography. **b.** The sensory response of an example putative L6CT neuron recorded in an awake NTSR1-cre mouse expressing ChR2-eYFP. **c.** Population PSTH overlaid with the z-scored sensory response of each putatitve L6CT neurons recorded in an awake mouse, thresholded at a firing rate of 200 (n = 44 neurons, 33 recordings, 7 mice). **d.** Same as ***c***, but for putative L6CT neurons recorded in mice under anesthesia (n = 24 neurons, 19 recordings, 4 mice). **e.** The mean +/- sem baseline-subtracted sensory response of the same neurons in ***c*** and ***d*** highlights that L6CT neurons in awake mice are more responsive than those recorded under anestheisa (p = 2.89e-2, determined using a one-sided Wilcoxon ranksum test). **f.** The mean +/- sem spontaneous activity of putative L6CT neurons recorded during wakefulness is not significantly different than that under anesthesia (p = 0.40, determined using a one-sided Wilcoxon

One important difference between our finding and the previously referenced reports on sparsely firing L6CT neurons is the state of the animal. To verify that the apparent responsiveness of L6CT neurons that we measured is related to the non-anesthetized/awake state of the animal as opposed to some other details of our recording and analysis, in a subset of recordings, we measured sensory response of L6CT neurons in a separate cohort of mice that were lightly anesthetized with isoflurane anesthesia (see Methods). Figure 2d shows the L6CT population sensory response in mice under anesthesia. We find that only 25% (6 out of 24) of putative L6CT neurons demonstrate a statistically significant change in spiking relative to pre-stimulus activity and the L6CT neuron baseline-subtracted population response is more than two times smaller than what we observe in the awake animal (Figure 2e). Importantly, anesthesia is known to suppress spontaneous cortical activity and alter the temporal structure of ongoing population activity (Constantinople & Bruno, 2011; Greenberg et al., 2008), which could bear an impact on the stimulus-evoked response computed here. But we find that there is no significant difference in spontaneous activity between L6CT neurons recorded in awake versus anesthetized mice (Figure 2f), indicating that the difference in L6CT neuron activity during wakefulness versus anesthesia is due to the responsiveness to the sensory stimulus and not drastic changes in baseline activity.

To further benchmark the L6CT sensory response, we compared it to that of all other simultaneously recorded RS neurons (excluding L6CT neurons). We find that other RS neurons exhibit an overall greater baseline-subtracted population sensory response compared to the response of the L6CT population (Figures 3a & 3b). Looking in more detail, the response of the other RS neurons in each individual layer is also greater than the response of L6CT neurons (Supplemental figure 1). Compared to the 53% of L6CT neurons that are sensory-responsive, 85% of other cortical RS neurons are sensory-responsive, consistent with previous reports (O’Connor et al., 2010; Pala & Stanley, 2022). Importantly, this difference in sensory response is not due to a systematic difference in spontaneous activity, as there is no significant difference in the spontaneous activity of L6CT neurons compared to other RS neurons (Figure 3c).

**Figure 3.**
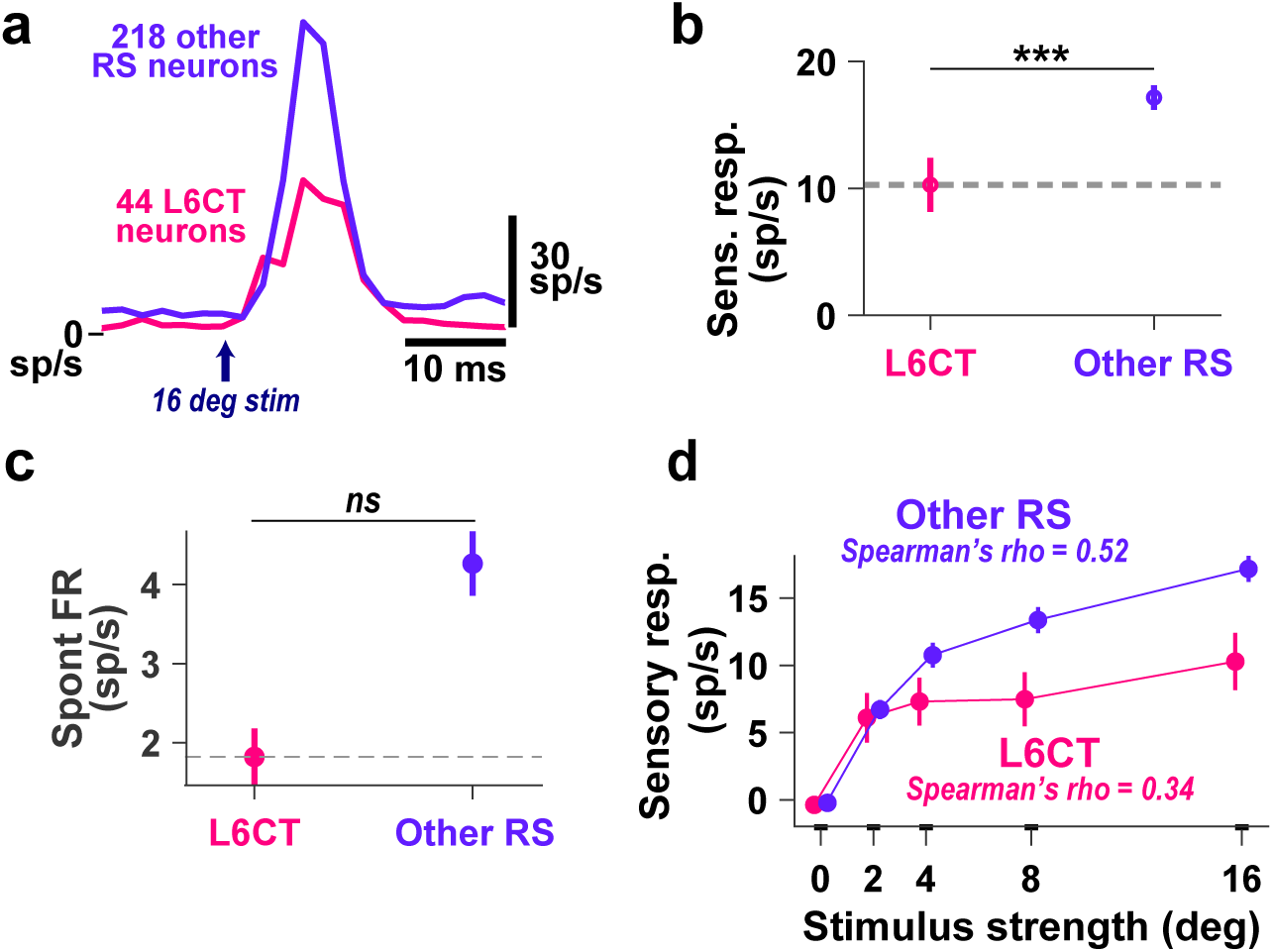
L6CT neurons are less sensory-responsive than other RS neurons during wakefulness. **a.** Grand PSTHs of the recorded L6CT population and all other RS neurons (excluding L6CT neurons) to a 16 deg sensory stimulus (n = 7 mice, 33 recordings). **b.** The mean +/- sem baseline-subtracted sensory response of the same neurons in ***a*** highlights that L6CT neurons in awake mice are less responsive than that of other RS neurons (p = 2.14e-4, determined using a one-sided Wilcoxon ranksum test). **c.** The mean +/- sem spontaneous firing rate of the same neurons in ***a*** is not significantly different between L6CT neurons and all other RS neurons (p = 0.40, determined using a two-sided Wilcoxon ranksum test). **d.** The baseline-subtracted stimulus response curve for the population of putatitve L6CT neurons and all other S1 RS neurons demonstrates that the mean +/- sem sensory response of L6CT neurons does not increase as much with increasing stimulus strength as that of other RS neurons.

We next characterized the sensitivity of L6CT neurons to stimulus strength compared to other RS neurons (Figure 3d). Here, the stimulus strengths are defined by their amplitude in degrees which corresponds to specific velocities, to which thalamocortical neurons are most sensitive (see Methods; Waiblinger et al., 2015). We observed that while the response of other cortical neurons continues to increase with increasing stimulus strength, this is not the case for L6CT neurons. Rather, after the initial increase to the 2 deg stimulus, the stimulus response curve for the population of L6CT neurons is relatively flat. The Spearman’s correlations computed from these curves highlights a weaker association between the population response and increasing stimulus strength for L6CT neurons compared to other RS neurons (L6CT neurons: rho = 0.34; Other RS neurons: rho = 0.52).

### L6CT neuron activity can better detect than discriminate sensory stimuli

The extent to which neurons encode a sensory stimulus is ultimately determined not only by the magnitude of the sensory response and the spontaneous activity, but also by the variability of the response across trials. So, we next quantified the extent to which the presence of the 16 deg sensory stimulus could be decoded from the activity of L6CT neurons (same data as in Figure 3) on an individual trial basis compared to other RS neurons. Figure 4a shows example probability density functions (pdf) fit using the spontaneous activity and stimulus-evoked activity (not baseline-subtracted in this case) for an example subset of 25 neurons from each cortical subpopulation. In these example subpopulation subsets, the pdfs for the L6CT neurons are similarly separable as compared to those for other RS neurons. Using bootstrapped pdf fits from each subpopulation, we decoded the presence or absence of the sensory stimulus using linear discriminant analysis (LDA; see Methods). We found that the signal versus noise distributions from the activity of L6CT neurons as well as that from the activity of other RS neurons were sufficiently separated such that the presence or absence of the stimulus can be reliably decoded above chance. Further, there was no significant improvement in decodability using the activity of other RS neurons compared to using the activity of L6CT neurons (Figure 4b).

**Figure 4.**
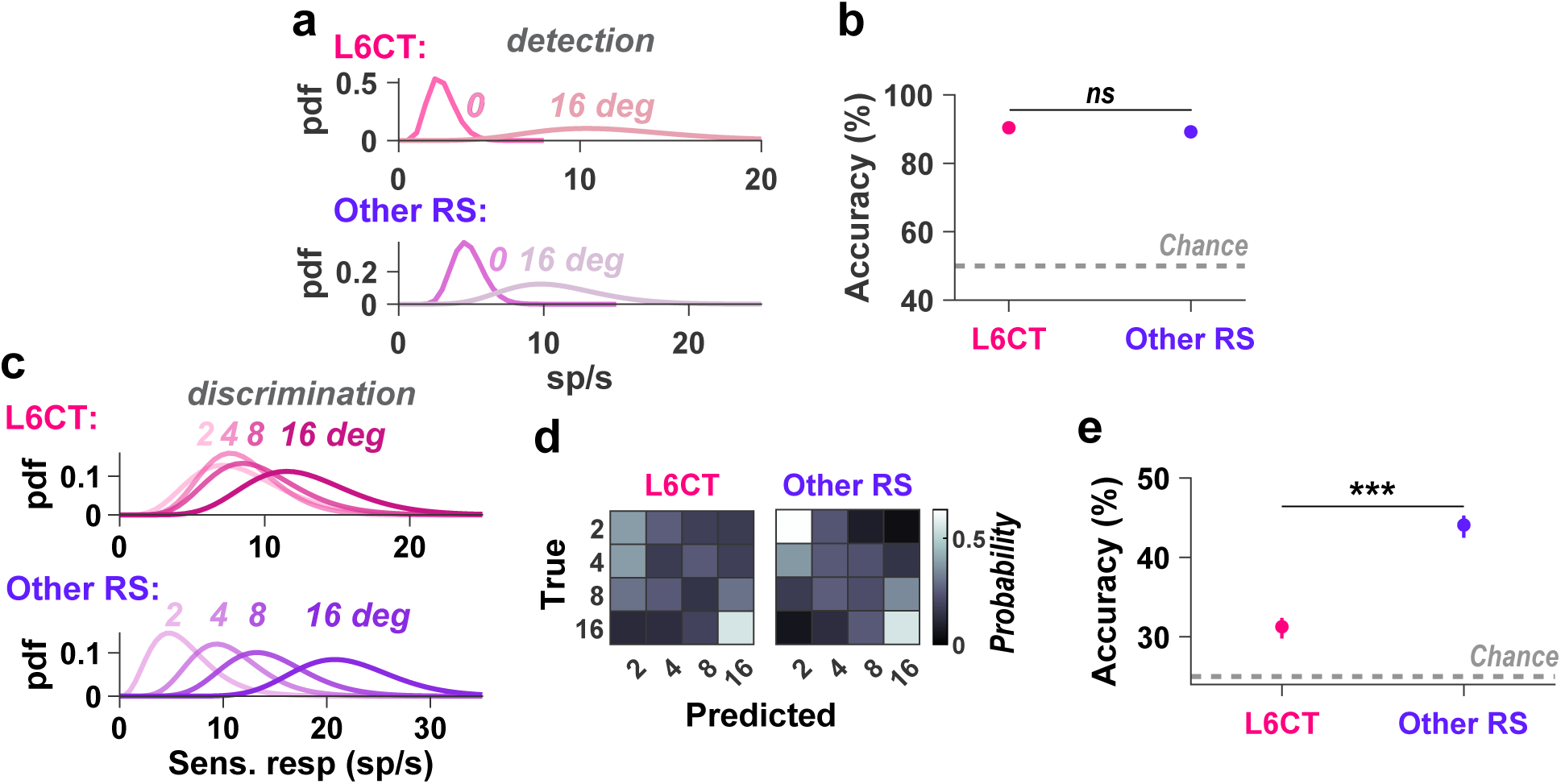
An ideal observer can reliably detect the presence of a sensory stimulus from the activity of L6CT neurons but the strength of the sensory stimulus is not as well discriminated. **a.** The parameterized probability density functions (pdfs) of the signal (sensory response to the 16 deg stimulus) and noise (spontaneous activity) across trials for an example subset of each recorded cortical subpopulation presented in. The pdfs are gamma distribution fits of the combined spiking of 25 neurons in 25 trials, both sampled with replacement. **b.** The mean accuracy of decoding the presence of a sensory stimulus from spontaneous activity across 1000 iterations of decoding tin each cortical subpopulation is not significantly different for L6CT neurons compared to other RS neurons (+/-99% confidence intervals; *** indicates p < 0.001). Decoding is performed using linear discriminant analysis on the spiking of 25 neurons in 25 trials. **c.** The parameterized probability density functions (pdfs) of the baseline-subtracted sensory response across trials for an example subset of each recorded cortical subpopulation. The pdfs are gamma distribution fits of the combined spiking of 25 neurons in 25 trials, both sampled with replacement. **d.** The confusion matrix indicating the discriminability of stimulus srength using neuronal activity of each cortical subpopulation. Decoding is performed using linear discriminant analysis on the spiking of 25 neurons in 25 trials. **e.** The mean accuracy from 1000 iterations of decoding the sensory stimulus from spontaneous activity using the spiking from each subpopulationof is lower for L6CT neurons compared to other RS neurons (+/- 99% confidence intervals; *** indicates p < 0.001; re-sampling neurons and trials with replacement).

To then assess the extent to which L6CT neurons encode the stimulus strength, we again fit pdfs, this time on the baseline-subtracted stimulus-evoked response across stimulus strength for each neuronal subpopulation. Figure 4c shows gamma distribution fit pdfs of the response across trials to each stimulus strength for an example subset of 25 neurons for each cortical subpopulation. In these example subpopulation subsets, the pdfs for the L6CT neurons are highly overlapped, whereas that of the other RS neurons are more separated.

Using the same method as that to decode the presence or absence of a stimulus, we then used bootstrapped spiking from each cortical subpopulation to decode the stimulus strength using LDA. The confusion matrix in Figure 4d shows the probability of the ideal observer predicting each stimulus strength given the true stimulus strength. Overall, we found that the stimulus strength can indeed be decoded above chance levels from L6CT population activity. However, the ideal observer performed significantly less well when decoding between stimulus strengths using the activity of L6CT neurons compared to when using the activity of other RS neurons (Figure 4d & e).

### L6CT neuron sensory responses reflect the behavioral outcome of mice performing in a sensory detection task

Our decoding analyses suggest that while L6CT neurons are not detailed encoders of the sensory stimulus strength compared to other S1 neurons, they do signal the presence of a sensory stimulus. This suggests a potential role for L6CT neurons in the detection of sensory stimuli, which we explored next. To study the engagement of L6CT neurons during sensory detection, we trained seven NTSR1-cre mice in a simple somatosensory detection task (Figure 5a). In this task, mice report the detection a whisker deflection by licking a spout for a water reward. Trials in which the animal accurately licks in response to a stimulus are referred to as “hit” trials, while “miss” trials refer to trials in which the animal does not respond despite the whisker being deflected. The response window within which licks were rewarded was one second long and licks occurring during the window lasting one second prior to the stimulus presentation resulted in the trial being aborted. The mice were trained using the 16 deg stimulus in addition to catch trials (0 deg stimulus) presented on 20% of trials. Once the mice learned the task, the difficulty of the task was pseudo-randomly varied across trials by presenting stimuli of varying strength: 0, 2, 4, 8, and 16 deg with equal probability (Figure 5a; see Methods; Ollerenshaw et al., 2012; Waiblinger et al., 2019). The mice learn to perform this task well and, upon learning, exhibit a monotonic relationship between the stimulus strength and the probability of the animal licking within the response window; that is that the animals accurately detected stronger stimuli more often than weaker stimuli (Figure 5b; Spearman’s rank correlation = 0.84; p = 1.7e-45). Additionally, the larger the stimulus strength, the faster the reaction time, as indicated by the negative monotonic relationship between the stimulus strength and the average lick latency, or the time to the first lick spout contact (Figure 5c). Across all non-zero strength sensory stimuli, the average lick latency across all sessions is 428 ms, consistent with prior reports of somatosensory detection tasks (Ollerenshaw et al., 2014; Stüttgen et al., 2006; Waiblinger et al., 2018, 2019, 2022).

**Figure 5.**
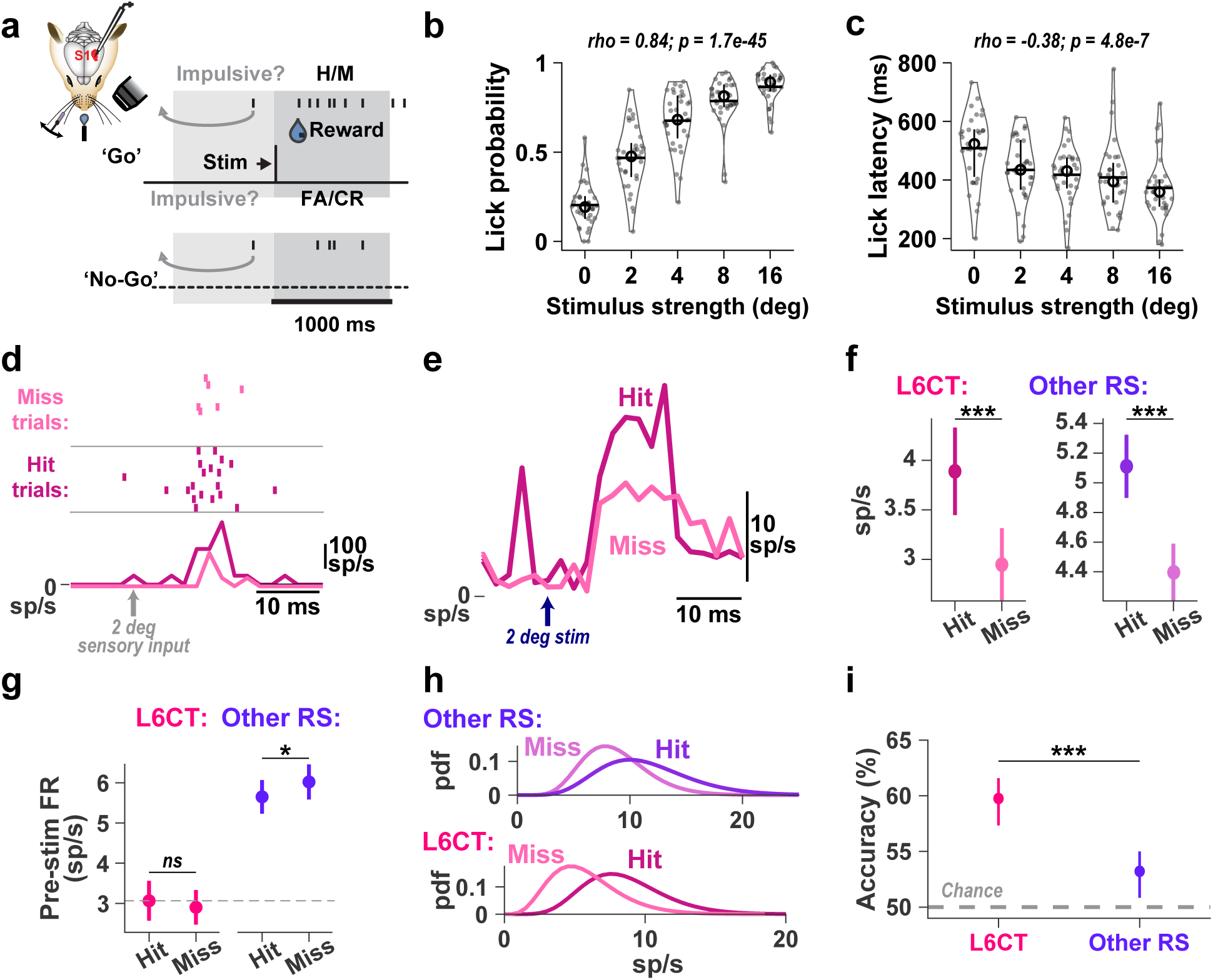
L6CT neuron sensory responses reflect the behavioral outcome. **a.** NTSR1-cre mice were trained to lick for a water reward within a one second response window after an individual whisker was precisely deflected to various strengths by varying the amplitude using a computer-controlled galvanometer. Any licks occurring within the second prior to the stimulus presentation resulted in the trial being aborted (see Methods). **b.** The probability of the trained mice licking within the response window increases monotonically with the stimulus strength (n = 33 behavioral sessions across the seven mice; for each violin plot, the circle indicates the median across sessions, the horizontal line indicates the mean, the vertical line indicates the range between the first and third quartiles, and the outline indicates the kernel density estimate of the lick probability across sessions; rho & p-value: Spearman’s rank correlation). **c.** Same as b, but for the latency of the first lick post-stimulus, which decreases with increasing stimulus strength. **d.** Rasters and PSTHs of an example L6CT neuron’s response to a 2 deg sensory stimulus parsed by the behavioral outcome. Trials where the animal successfully licked in the response window after the stimulus are hit trials and miss trials are trials when the animal did not lick despite the presentation of the stimulus. **e.** Grand PSTHs of the recorded L6CT population (n = 44 neurons) to the 2 deg sensory stimulus, parsed by the behavioral outcome. Note that the transient and large increase in spiking occurring prior to the stimulus onset in hit trials is from a single neuron; for this neuron’s recording, there is only one hit trial, in which the neuron fired only one spike, which results in a high firing rate that dominates the mean across neurons for that bin. **f.**The mean +/- sem baseline-subtracted sensory response of the populations of L6CT and other RS neurons is larger in hit trials than in miss trials. **g.** The mean +/- sem pre-stimulus firing rate (500 ms prior to the sensory stimulus) in hit vs miss trials is unchanged in L6CT neurons, whereas the pre-stimulus spiking in other RS neurons is significantly higher in miss trials. **h.** The parameterized probability density functions (pdfs) of the 2 deg sensory response across trials for an example subset of each cortical subpopulation in d are better separated for L6CT neurons compared to other RS neurons. Pdfs are gamma distribution fits of the combined spiking of 25 neurons in 25 trials, both sampled with replacement. **i.** The mean accuracy from 1000 iterations of decoding the animal’s behavioral performance using the activity of a subset of neurons from each cortical subpopulation in d is higher for L6CT neurons compared to that for other RS neurons (+/- 99% confidence intervals; *** indicates p < 0.001; re-sampling neurons and trials with replacement). Decoding is performed using linear discriminant analysis on the spiking of 25 neurons in 25 trials with the 2 deg stimulus.

We performed silicon probe recordings of populations of individual neurons throughout S1, including optotagged L6CT neurons as presented in Figure 1, across 33 sessions in the seven trained mice. To determine the extent to which L6CT activity relates to the animal’s behavior, we parsed the activity of L6CT neurons based on the behavioral outcome. Here and in subsequent analyses, we present the response to the 2 deg stimulus which is the behavioral response threshold stimulus, or the stimulus that the animals accurately detect in approximately 50% of trials on average across sessions. Use of the threshold stimulus enables roughly equivalent numbers of hit and miss trials for subsequent analyses.

Figure 5d shows the response of an example L6CT neuron to the behavioral response threshold stimulus parsed by the behavioral outcome. This neuron elicits more spikes to the same stimulus in hit trials compared to miss trials. This greater response in hit trials is consistent with the population of recorded L6CT neurons (Figure 5e). This is also consistent with what we (Figure 5f) and others (Kyriakatos et al., 2017; Speed et al., 2019) observe for other RS neurons.

In addition to the sensory response, the ongoing cortical activity also plays an important role in sensory perception and it has previously been observed in similarly simple detection tasks that the behavioral outcome is correlated with differences in the ongoing firing rate of cortical neurons (McGinley et al., 2015; Speed et al., 2019). To assess the extent to which the difference in sensory response is based on changes in ongoing activity, we quantified the pre-stimulus firing rate of L6CT neurons. Contrary to the pre-stimulus firing rate of other RS neurons which is slightly higher in miss compared to hit trials, which is consistent with other reports (Waiblinger et al., 2018), the pre-stimulus firing rate of L6CT neurons is unchanged between hit and miss trials (Figure 5g).

Given the observed difference in neural activity between hit and miss trials, we assessed the accuracy of predicting the behavioral outcome of the animal on a single trial basis using the spiking activity of each cortical subpopulation. Again, for 1000 iterations, we sub-sampled with replacement the stimulus-evoked spiking in 25 trials of each behavioral outcome for 25 neurons from each cortical subpopulation. Figure 5h shows the fit pdfs from an example iteration for each cortical subpopulation. In this example parameterized subset of neuronal responses, given both the change in mean as well as the variance of the distributions, it is clear that the hit and miss trial distributions for L6CT neurons are more separable compared to that of the other RS neurons. Indeed, when feeding the across-trial spiking subsets into an LDA decoder, we observe that while the behavioral outcome can be decoded above chance levels for both subpopulations on average across all iterations, the spiking from L6CT neurons can significantly better decode the behavioral outcome of the animal (Figure 5i).

## Discussion

Although there are other distinct corticothalamic pathways that have been described (Kirchgessner et al., 2021; Usrey & Sherman, 2018, 2019), the predominant pathway is from pyramidal neurons in cortical layer 6 (L6CT) that project and provide modulatory input to the thalamus (Guillery & Sherman, 2002; Sherman, 2016). Outnumbering ascending projections to the thalamus in sensory pathways nearly 10:1 (Deschênes et al., 1998; Sherman & Koch, 1986), corticothalamic feedback has long been hypothesized to serve a critical role in sensory signaling, providing a potential mechanistic underpinning of the dynamic gating of information flow from the sensory periphery to higher cognitive centers (Sherman & Guillery, 1998). Yet these pathways have traditionally been difficult to study in detail due to the complexity of the circuits in which they are embedded. The advent of tools to precisely identify L6CT neurons for targeted measurement in-vivo (Gong et al., 2003, 2007) has generated significant interest towards unveiling the function of L6CT neurons in a growing number of studies (Born et al., 2021; Bortone et al., 2014; Denman & Contreras, 2015; Dimwamwa et al., 2024; Folkard et al., 2026; Hasse & Briggs, 2017; Kirchgessner et al., 2020, 2021; Olsen et al., 2012; Pauzin et al., 2019; Pauzin & Krieger, 2018;

Reinhold et al., 2023; Russo et al., 2025; Spacek et al., 2022; Ziegler et al., 2023). But studies performing direct measurements of L6CT activity during wakefulness are rare (but see Augustinaite & Kuhn, 2020; Briggs & Usrey, 2009; Dash et al., 2022; Swadlow, 1987, 1989; Swadlow & Hicks, 1996), and even less so in behaving animals (but see Clayton et al., 2021; Oliveira & Kanold, 2026; Vélez-Fort et al., 2018). To further complicate matters, in addition to receiving direct, bottom-up, monosynaptic input from first order thalamus, L6CT neurons also receive top-down intracortical synaptic inputs that positions them as modulators of thalamic gating (Clayton et al., 2021; W. Guo et al., 2017; Whilden et al., 2021), suggesting that the relative role of L6CT neurons could depend strongly on the timescale in question. So while our understanding about these neurons is growing, we are left without context as to *when*, exactly, L6CT neurons would exert their modulatory function in sensory signaling and behavior. In this study, we provide a glimpse of the contexts in which L6CT neurons may modulate thalamocortical activity by determining that L6CT neurons are not as inactive and functionally sparse as previously shown in both their stimulus-evoked and spontaneous spiking in awake and behaving animals. More specifically, we demonstrate that optogenetically identified L6CT neurons in the awake animal do indeed convey information from the sensory periphery, contrary to the activity of L6CT neurons measured under isoflurane anesthesia as in a number of studies. Intriguingly, unlike other cortical excitatory neurons, L6CT neuron activity is reflective of the presence or absence of a stimulus, but not the stimulus strength. This suggests that in this context, L6CT neurons may primarily support the detection of sensory inputs rather than the discrimination of fine detail. This prompted the measuring of the activity of L6CT neurons in mice performing a simple detection task to reveal the engagement of L6CT neurons in the detection of somatosensory inputs, where we demonstrate that they robustly encode the behavioral outcome. In the larger context of the complex circuit, this activity would in turn modulate L6CT neurons’ thalamic and cortical targets (Dimwamwa et al., 2024; Russo et al., 2025), which could have longer timescale effects on sensory perception.

Despite the technical challenges, there is growing evidence for L6CT neurons playing a role in bottom-up sensory processing (Andolina et al., 2007; Born et al., 2021; Briggs & Usrey, 2009; Denman & Contreras, 2015; Hasse & Briggs, 2017; Mease et al., 2014; Olsen et al., 2012; Temereanca et al., 2008; Temereanca & Simons, 2004) as well as growing evidence for L6CT neurons playing top-down and extra-sensory roles, such as in brain state-related modulation of sensory cortex (Reinhold et al., 2023) and motor-related modulation of sensory processing (Augustinaite & Kuhn, 2020; Clayton et al., 2021; Dash et al., 2022). In a previous study, we found that optogenetically driven L6CT activity had the overall effect of increasing cortical excitability across cortical layers. Under these same conditions, L6CT neurons bi-directionally modulated first-order thalamic neurons such that high levels of L6CT firing rate and synchrony increased excitability of thalamic neurons while low levels had an opposite, suppressive effect (Dimwamwa et al., 2024). Based on those findings, the general levels of L6CT activity observed in this study would likely in turn increase cortical excitability across layers and decrease the excitability of first order thalamic relay neurons, although this was not measured directly here. More recently, we found that optogenetic activation of L6CT neurons induces high frequency oscillations in the cortical LFP (Russo et al., 2025), further placing the L6CT neurons in the potential role of dynamic mediator of cortical excitability. To fully elucidate the role of the L6CT neurons as both recipients and modulators of first order thalamic and intracortical projections, further studies are needed that directly investigate the causal role of these neurons in behavior.

Despite the sensory input being exactly the same in hit versus miss trials, we find that cortical responses are larger in hit versus miss trials for both the L6CT subpopulation as well as for the subpopulation of other RS neurons. Consistent with previous reports (Kyriakatos et al., 2017; Speed et al., 2019), this finding reiterates the notion that sensory processing involves information flow of inputs from the periphery through the processing hierarchy combined with top-down and neuromodulator influences to shape the neuronal representation of the sensory inputs (Castro-Alamancos, 2004; Gilbert & Li, 2013). In the presented data, we excluded trials with whisker movements and also demonstrate that the animal licks, on average, >400 ms after the sensory stimulus, well beyond our analysis window. So what seems likely here is that the difference in hit versus miss trials reflects different levels of excitability in cortex, the source of which must be disentangled in future studies. Consistent with this, beyond just the L6CT neurons, our results here show that the pre-stimulus firing rate and sensory response of primary sensory cortical neurons reflect the behavior of the animal. This is consistent with observations of layer 2/3 neurons in a similar detection task (Yang et al., 2016) as well measurements from V1 in a visual detection task (Speed et al., 2019). However, it is important to note that Sachidhanandam et al. reported that the pre-stimulus and early components of the stimulus-evoked response are unchanged between hit and miss trials in S1 during a somatosensory detection task (Sachidhanandam et al., 2013). Relatedly, S1 neurons in primates do not show behavior-related signaling (de Lafuente & Romo, 2005, 2006). While one explanation for the difference could be that the results presented here are for threshold-level stimuli, we find that the behavioral outcome of the animal relates to cortical activity across all tested stimulus strengths (data not shown). Thus, a more likely explanation is that our results sample neurons from all layers, with a high representation of neurons from layer 5, whereas Sachidhanandam et al. report on neurons exclusively from layers 2/3. Indeed, layer-specific differences are known (Speed et al., 2019). More generally, the exact details of the task can also influence the role of primary cortical neurons (Hong et al., 2018; Park et al., 2022; Rodgers et al., 2021). See Waiblinger et al., 2026 for a more extensive discussion on this topic (Waiblinger et al., 2026).

Taken together, we reveal that L6CT neurons convey information concerning the bottom-up inputs, the timing of which is fast enough to contribute to the sensory response of neurons throughout the thalamocortical circuit. However, the L6CT activity did not scale with stimulus strength whereas the performance of the animal did scale with stimulus strength, suggesting L6CT neurons are not a principal driver of the behavior. We demonstrate that L6CT neuron activity relates to whether it was a hit or miss trial, therefore suggesting a broader role for L6CT neurons in the gating of a given sensory stimulus. Further, the results of our study provide a specific range of activity patterns in which L6CT neurons operate during sensory perception, contextualizing previous manipulation studies (Born et al., 2021; Bortone et al., 2014; Denman & Contreras, 2015; Dimwamwa et al., 2024; Folkard et al., 2026; Hasse & Briggs, 2017; Kirchgessner et al., 2020, 2021; Olsen et al., 2012; Pauzin et al., 2019; Pauzin & Krieger, 2018; Reinhold et al., 2023; Russo et al., 2025; Spacek et al., 2022; Ziegler et al., 2023). Although the findings here were obtained in experiments in the somatosensory system, and specifically the vibrissa pathway, these fundamental results likely generalize to other sensory pathways. Indeed, L6CT neurons in the primary visual (V1) and auditory (A1) cortices are also sensory-responsive during wakefulness (Augustinaite & Kuhn, 2020; Cardoso de Oliveira & Kanold, 2026; Clayton et al., 2021). We build on these results and show that what differentiates the sensory responses of L6CT neurons from other cortical neurons is their sensitivity to the strength of the bottom-up inputs. The lack of strength sensitivity suggests a role of L6CT neurons in signaling the presence of sensory inputs, rather than in encoding the fine details, potentially tying them to gating in tactile detection, one of the most fundamental functional roles of the pathway.

## Contributions

Elaida Dimwamwa and Garrett B. Stanley conceptualized the project and designed the study. Elaida Dimwamwa, Christian Waiblinger, and Nelson H. Chang carried out all experiments. Elaida Dimwamwa analyzed the data with contributions from Nelson H. Chang. Elaida Dimwamwa and Garrett B. Stanley wrote the paper; all authors edited the paper. Elaida Dimwamwa and Garrett B. Stanley acquired funding.

## Acknowledgements

This work was supported by NIH National Institute of Neurological Disorders and Stroke (NINDS) BRAIN Grant R01NS104928 and NINDS BRAIN Grant RF1NS128896. EDD was supported by a National Science Foundation Graduate Research Fellowship and the Howard Hughes Medical Institute through the James H. Gilliam Fellowships for Advanced Study program. We would like to thank David A. Weiss for assistance in the analysis.

---

**Figure S1.**
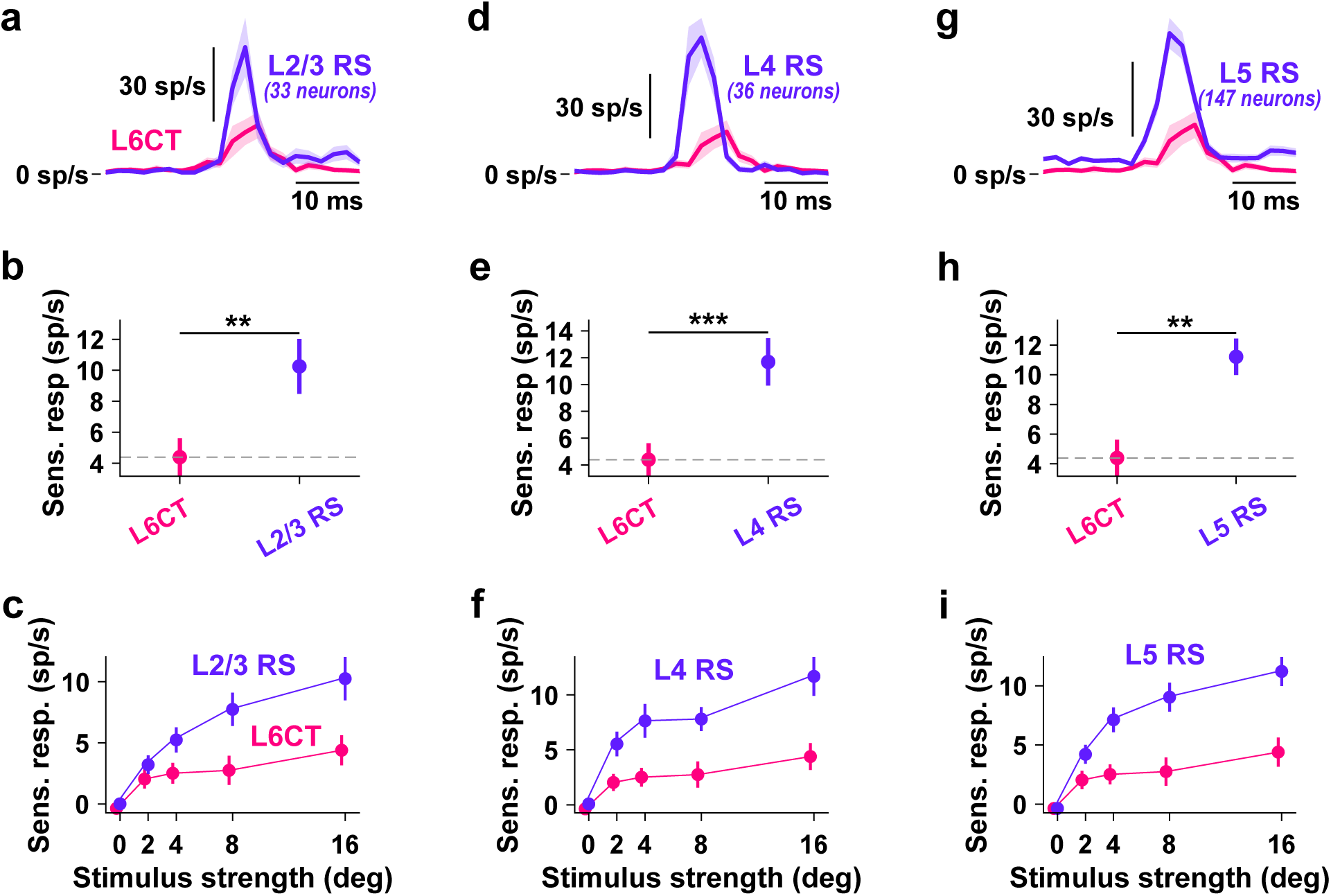
L6CT neurons are less sensory-responsive than other RS neurons in other layers during wakefulness. **a.** Grand PSTHs of the recorded L6CT population and L2/3 RS neurons to a 16 deg sensory stimulus. **b.** The mean +/- sem baseline-subtracted sensory response of the same neurons in ***a*** highlights that L6CT neurons in awake mice are less responsive than L2/3 RS neurons. **c.** The baseline-subtracted stimulus response curve for the population of putatitve L6CT neurons and L2/3 RS neurons demonstrates that the mean +/- sem sensory response of L6CT neurons does not increase as much with increasing stimulus strength as that of L2/3 RS neurons. **d-f.** Same as a-c, butfor L4 RS neurons. **g-i.** Same as a-c, but for L5 RS neurons. Note that L6 neurons are not shown because there are insufficient numbers of non-L6CT L6 neurons.

## References

Andolina, I. M., Jones, H. E., Wang, W., & Sillito, A. M. (2007). Corticothalamic feedback enhances stimulus response precision in the visual system. Proceedings of the National Academy of Sciences, 104(5), 1685–1690. 10.1073/pnas.0609318104

Augustinaite, S., & Kuhn, B. (2020). Complementary Ca2+ Activity of Sensory Activated and Suppressed Layer 6 Corticothalamic Neurons Reflects Behavioral State. Current Biology, 30(20), 3945–3960.e5. 10.1016/j.cub.2020.07.069

Barthó, P., Hirase, H., Monconduit, L., Zugaro, M., Harris, K. D., & Buzsáki, G. (2004). Characterization of Neocortical Principal Cells and Interneurons by Network Interactions and Extracellular Features. Journal of Neurophysiology, 92(1), 600–608. 10.1152/jn.01170.2003

Bolus, M. F., Willats, A. A., Whitmire, C. J., Rozell, C. J., & Stanley, G. B. (2018). Design strategies for dynamic closed-loop optogenetic neurocontrol *in vivo*. Journal of Neural Engineering, 15(2), 026011. 10.1088/1741-2552/aaa506

Borden, P. Y., Wright, N. C., Morrissette, A. E., Jaeger, D., Haider, B., & Stanley, G. B. (2022). Thalamic bursting and the role of timing and synchrony in thalamocortical signaling in the awake mouse. Neuron, 110(17), 2836–2853.e8. 10.1016/j.neuron.2022.06.008

Born, G., Schneider-Soupiadis, F. A., Erisken, S., Vaiceliunaite, A., Lao, C. L., Mobarhan, M. H., Spacek, M. A., Einevoll, G. T., & Busse, L. (2021). Corticothalamic feedback sculpts visual spatial integration in mouse thalamus. Nature Neuroscience, 24(12), 1711–1720. 10.1038/s41593-021-00943-0

Bortone, D. S., Olsen, S. R., & Scanziani, M. (2014). Translaminar Inhibitory Cells Recruited by Layer 6 Corticothalamic Neurons Suppress Visual Cortex. Neuron, 82(2), 474–485. 10.1016/j.neuron.2014.02.021

Briggs, F., & Usrey, W. M. (2009). Parallel Processing in the Corticogeniculate Pathway of the Macaque Monkey. Neuron, 62(1), 135–146. 10.1016/j.neuron.2009.02.024

Carandini, M., & Churchland, A. K. (2013). Probing perceptual decisions in rodents. Nature Neuroscience, 16(7), 824–831. 10.1038/nn.3410

Cardoso de Oliveira, M., & Kanold, P. O. (2026). Layer 6 corticothalamic neurons show diverse and dynamic responses that support a role in cortical gain control in noisy environments (p. 2026.05.22.726967). bioRxiv. 10.64898/2026.05.22.726967

Castro-Alamancos, M. A. (2004). Dynamics of sensory thalamocortical synaptic networks during information processing states. Progress in Neurobiology, 74(4), 213–247. 10.1016/j.pneurobio.2004.09.002

Clayton, K. K., Williamson, R. S., Hancock, K. E., Tasaka, G., Mizrahi, A., Hackett, T. A., & Polley, D. B. (2021). Auditory Corticothalamic Neurons Are Recruited by Motor Preparatory Inputs. Current Biology, 31(2), 310–321.e5. 10.1016/j.cub.2020.10.027

Constantinople, C. M., & Bruno, R. M. (2011). Effects and Mechanisms of Wakefulness on Local Cortical Networks. Neuron, 69(6), 1061–1068. 10.1016/j.neuron.2011.02.040

Constantinople, C. M., & Bruno, R. M. (2013). Deep Cortical Layers Are Activated Directly by Thalamus. Science, 340(6140), 1591–1594. 10.1126/science.1236425

Crandall, S. R., Cruikshank, S. J., & Connors, B. W. (2015). A Corticothalamic Switch: Controlling the Thalamus with Dynamic Synapses. Neuron, 86(3), 768–782. 10.1016/j.neuron.2015.03.040

Cudeiro, J., Rivadulla, C., & Grieve, K. L. (2000). Visual response augmentation in cat (and macaque) LGN: potentiation by corticofugally mediated gain control in the temporal domain. European Journal of Neuroscience.

Dash, S., Autio, D. M., & Crandall, S. R. (2022). State-Dependent Modulation of Activity in Distinct Layer 6 Corticothalamic Neurons in Barrel Cortex of Awake Mice. The Journal of Neuroscience, 42(34), 6551–6565. 10.1523/JNEUROSCI.2219-21.2022

de Lafuente, V., & Romo, R. (2005). Neuronal correlates of subjective sensory experience. Nature Neuroscience, 8(12), 1698–1703. 10.1038/nn1587

de Lafuente, V., & Romo, R. (2006). Neural correlate of subjective sensory experience gradually builds up across cortical areas. Proceedings of the National Academy of Sciences of the United States of America, 103(39), 14266–14271. 10.1073/pnas.0605826103

Denman, D. J., & Contreras, D. (2015). Complex Effects on In Vivo Visual Responses by Specific Projections from Mouse Cortical Layer 6 to Dorsal Lateral Geniculate Nucleus. Journal of Neuroscience, 35(25), 9265–9280. 10.1523/JNEUROSCI.0027-15.2015

Deschênes, M., Veinante, P., & Zhang, Z.-W. (1998). The organization of corticothalamic projections: Reciprocity versus parity. Brain Research Reviews, 28(3), 286–308. 10.1016/S0165-0173(98)00017-4

DiCarlo, J. J., Lane, J. W., Hsiao, S. S., & Johnson, K. O. (1996). Marking microelectrode penetrations with fluoresce& dyes. Journal of Neuroscience Methods, 64, 75–81. 10.1016/0165-0270(95)00113-1

Dimwamwa, E. D., Pala, A., Chundru, V., Wright, N. C., & Stanley, G. B. (2024). Dynamic corticothalamic modulation of the somatosensory thalamocortical circuit during wakefulness. Nature Communications, 15(1), 3529. 10.1038/s41467-024-47863-8

Eyding, D., Macklis, J. D., Neubacher, U., Funke, K., & Wörgötter, F. (2003). Selective Elimination of Corticogeniculate Feedback Abolishes the Electroencephalogram Dependence of Primary Visual Cortical Receptive Fields and Reduces Their Spatial Specificity. The Journal of Neuroscience, 23(18), 7021–7033. 10.1523/JNEUROSCI.23-18-07021.2003

Folkard, R., Isaías-Camacho, E. U., & Groh, A. (2026). Corticothalamic layer 6 controls cortical activity and thalamic firing mode in a bidirectional manner. Cerebral Cortex, 36(3), bhag029. 10.1093/cercor/bhag029

Gilbert, C. D., & Li, W. (2013). Top-down influences on visual processing. Nature Reviews Neuroscience, 14(5), 350–363. 10.1038/nrn3476

Gong, S., Doughty, M., Harbaugh, C. R., Cummins, A., Hatten, M. E., Heintz, N., & Gerfen, C. R. (2007). Targeting Cre Recombinase to Specific Neuron Populations with Bacterial Artificial Chromosome Constructs. Journal of Neuroscience, 27(37), 9817–9823. 10.1523/JNEUROSCI.2707-07.2007

Gong, S., Zheng, C., Doughty, M. L., Losos, K., Didkovsky, N., Schambra, U. B., Nowak, N. J., Joyner, A., Leblanc, G., Hatten, M. E., & Heintz, N. (2003). A gene expression atlas of the central nervous system based on bacterial artificial chromosomes. Nature, 425(6961), 917–925. 10.1038/nature02033

Greenberg, D. S., Houweling, A. R., & Kerr, J. N. D. (2008). Population imaging of ongoing neuronal activity in the visual cortex of awake rats. Nature Neuroscience, 11(7), 749– 751. 10.1038/nn.2140

Guillery, R. W., & Sherman, S. M. (2002). Thalamic Relay Functions and Their Role in Corticocortical Communication: Generalizations from the Visual System. 33, 163–175. 10.1016/S0896-6273(01)00582-7

Guo, W., Clause, A. R., Barth-Maron, A., & Polley, D. B. (2017). A Corticothalamic Circuit for Dynamic Switching between Feature Detection and Discrimination. Neuron, 95(1), 180–194.e5. 10.1016/j.neuron.2017.05.019

Guo, Z. V., Hires, S. A., Li, N., O’Connor, D. H., Komiyama, T., Ophir, E., Huber, D., Bonardi, C., Morandell, K., Gutnisky, D., Peron, S., Xu, N., Cox, J., & Svoboda, K. (2014). Procedures for Behavioral Experiments in Head-Fixed Mice. PLoS ONE, 9(2), e88678. 10.1371/journal.pone.0088678

Hasse, J. M., & Briggs, F. (2017). Corticogeniculate feedback sharpens the temporal precision and spatial resolution of visual signals in the ferret. Proceedings of the National Academy of Sciences, 114(30), E6222–E6230. 10.1073/pnas.1704524114

Hong, Y. K., Lacefield, C. O., Rodgers, C. C., & Bruno, R. M. (2018). Sensation, movement and learning in the absence of barrel cortex. Nature, 561(7724), 542–546. 10.1038/s41586-018-0527-y

Hooks, B. M., Hires, S. A., Zhang, Y.-X., Huber, D., Petreanu, L., Svoboda, K., & Shepherd, G. M. G. (2011). Laminar Analysis of Excitatory Local Circuits in Vibrissal Motor and Sensory Cortical Areas. PLoS Biology, 9(1), e1000572. 10.1371/journal.pbio.1000572

Kirchgessner, M. A., Franklin, A. D., & Callaway, E. M. (2020). Context-dependent and dynamic functional influence of corticothalamic pathways to first- and higher-order visual thalamus. Proceedings of the National Academy of Sciences, 117(23), 13066–13077. 10.1073/pnas.2002080117

Kirchgessner, M. A., Franklin, A. D., & Callaway, E. M. (2021). Distinct “driving” versus “modulatory” influences of different visual corticothalamic pathways. Current Biology, 31(23), 5121–5137.e7. 10.1016/j.cub.2021.09.025

Kvitsiani, D., Ranade, S., Hangya, B., Taniguchi, H., Huang, J. Z., & Kepecs, A. (2013). Distinct behavioural and network correlates of two interneuron types in prefrontal cortex. Nature, 498(7454), 363–366. 10.1038/nature12176

Kwegyir-Afful, E. E., & Simons, D. J. (2009). Subthreshold Receptive Field Properties Distinguish Different Classes of Corticothalamic Neurons in the Somatosensory System. Journal of Neuroscience, 29(4), 964–972. 10.1523/JNEUROSCI.3924-08.2009

Kyriakatos, A., Sadashivaiah, V., Zhang, Y., Motta, A., Auffret, M., & Petersen, C. C. H. (2017). Voltage-sensitive dye imaging of mouse neocortex during a whisker detection task. Neurophotonics, 4(3), 031204. 10.1117/1.NPh.4.3.031204

Lee, S., Carvell, G. E., & Simons, D. J. (2008). Motor modulation of afferent somatosensory circuits. Nature Neuroscience, 11(12), 1430–1438. 10.1038/nn.2227

Liew, Y. J., Dimwamwa, E. D., Wright, N. C., Zhang, Y., & Stanley, G. B. (2025). Multiple Distinct Timescales of Rapid Sensory Adaption in the Thalamocortical Circuit. The Journal of Neuroscience, 45(14), e1057242025. 10.1523/JNEUROSCI.1057-24.2025

Liew, Y. J., Pala, A., Whitmire, C. J., Stoy, W. A., Forest, C. R., & Stanley, G. B. (2021). Inferring thalamocortical monosynaptic connectivity in vivo. Journal of Neurophysiology, 125(6), 2408–2431. 10.1152/jn.00591.2020

Masino, S. A., Kwon, M. C., Dory, Y., & Frostig, R. D. (1993). Characterization of functional organization within rat barrel cortex using intrinsic signal optical imaging through a thinned skull. Proceedings of the National Academy of Sciences, 90(21), 9998–10002. 10.1073/pnas.90.21.9998

McCormick, D. A., & von Krosigk, M. (1992). Corticothalamic activation modulates thalamic firing through glutamate “metabotropic” receptors. Proceedings of the National Academy of Sciences, 89(7), 2774–2778. 10.1073/pnas.89.7.2774

McGinley, M. J., David, S. V., & McCormick, D. A. (2015). Cortical Membrane Potential Signature of Optimal States for Sensory Signal Detection. 87(1), 179–192. 10.1016/j.neuron.2015.05.038

Mease, R. A., Krieger, P., & Groh, A. (2014). Cortical control of adaptation and sensory relay mode in the thalamus. Proceedings of the National Academy of Sciences, 111(18), 6798–6803. 10.1073/pnas.1318665111

O’Connor, D. H., Clack, N. G., Huber, D., Komiyama, T., Myers, E. W., & Svoboda, K. (2010). Vibrissa-Based Object Localization in Head-Fixed Mice. The Journal of Neuroscience, 30(5), 1947–1967. 10.1523/JNEUROSCI.3762-09.2010

Ollerenshaw, D. R., Bari, B. A., Millard, D. C., Orr, L. E., Wang, Q., & Stanley, G. B. (2012). Detection of tactile inputs in the rat vibrissa pathway. Journal of Neurophysiology, 108(2), 479–490. 10.1152/jn.00004.2012

Ollerenshaw, D. R., Zheng, H. J. V., Millard, D. C., Wang, Q., & Stanley, G. B. (2014). The adaptive trade-off between detection and discrimination in cortical representations and behavior. Neuron, 81(5), 1152–1164. 10.1016/j.neuron.2014.01.025

Olsen, S. R., Bortone, D. S., Adesnik, H., & Scanziani, M. (2012). Gain control by layer six in cortical circuits of vision. Nature, 483(7387), 47–52. 10.1038/nature10835

Pachitariu, M., Sridhar, S., Pennington, J., & Stringer, C. (2024). Spike sorting with Kilosort4 | Nature Methods. 10.1038/s41592-024-02232-7

Pala, A., & Stanley, G. B. (2022). Ipsilateral Stimulus Encoding in Primary and Secondary Somatosensory Cortex of Awake Mice. The Journal of Neuroscience, 42(13), 2701– 2715. 10.1523/JNEUROSCI.1417-21.2022

Park, J. M., Hong, Y. K., Rodgers, C. C., Dahan, J. B., Harano, N., Schmidt, E. R., & Bruno, R. M. (2022). Deep and superficial layers of the primary somatosensory cortex are critical for whisker-based texture discrimination in mice (p. 2020.08.12.245381). bioRxiv. 10.1101/2020.08.12.245381

Pauzin, F. P., & Krieger, P. (2018). A Corticothalamic Circuit for Refining Tactile Encoding. Cell Reports, 23(5), 1314–1325. 10.1016/j.celrep.2018.03.128

Pauzin, F. P., Schwarz, N., & Krieger, P. (2019). Activation of Corticothalamic Layer 6 Cells Decreases Angular Tuning in Mouse Barrel Cortex. Frontiers in Neural Circuits, 13, 67. 10.3389/fncir.2019.00067

Ramón Y Cajal, S. (1906). The structure and connexions of neurons. Nobel Lecture.

Reinhold, K., Resulaj, A., & Scanziani, M. (2023). Brain State-Dependent Modulation of Thalamic Visual Processing by Cortico-Thalamic Feedback. The Journal of Neuroscience, 43(9), 1540–1554. 10.1523/JNEUROSCI.2124-21.2022

Rodgers, C. C., Nogueira, R., Pil, B. C., Greeman, E. A., Park, J. M., Hong, Y. K., Fusi, S., & Bruno, R. M. (2021). Sensorimotor strategies and neuronal representations for shape discrimination. Neuron, 109(14), 2308–2325.e10. 10.1016/j.neuron.2021.05.019

Rossant, C., & Harris, K. D. (2013). Hardware-accelerated interactive data visualization for neuroscience in Python. Frontiers in Neuroinformatics, 7. 10.3389/fninf.2013.00036

Russo, S., Dimwamwa, E. D., & Stanley, G. B. (2025). Layer 6 Corticothalamic Neurons Induce High Gamma Oscillations Through Cortico-cortical and Cortico-thalamo-cortical Pathways. Journal of Neuroscience, 45(42). 10.1523/JNEUROSCI.0094-25.2025

Sachidhanandam, S., Sreenivasan, V., Kyriakatos, A., Kremer, Y., & Petersen, C. C. H. (2013). Membrane potential correlates of sensory perception in mouse barrel cortex. Nature Neuroscience, 16(11), 1671–1677. 10.1038/nn.3532

Sederberg, A. J., Pala, A., Zheng, H. J. V., He, B. J., & Stanley, G. B. (2019). State-aware detection of sensory stimuli in the cortex of the awake mouse. PLOS Computational Biology, 15(5), e1006716. 10.1371/journal.pcbi.1006716

Sherman, S. M. (2016). Thalamus plays a central role in ongoing cortical functioning. Nature Neuroscience, 19(4), 533–541. 10.1038/nn.4269

Sherman, S. M., & Guillery, R. W. (1998). On the actions that one nerve cell can have on another: Distinguishing “drivers” from “modulators.” Proceedings of the National Academy of Sciences, 95(12), 7121–7126. 10.1073/pnas.95.12.7121

Sherman, S. M., & Koch, C. (1986). The control of retinogeniculate transmission in the mammalian lateral geniculate nucleus. Experimental Brain Research, 63(1). 10.1007/BF00235642

Sillito, A. M., Jones, H. E., Gerstein, George L., & West, D. C. (1994). Feature-linked synchronization of thalamic relay cell firing induced by feedback from visual cortex. 369, 479–482. 10.1038/369479a0

Sofroniew, N. J., Vlasov, Y. A., Hires, S. A., Freeman, J., & Svoboda, K. (2015). Neural coding in barrel cortex during whisker-guided locomotion. eLife, 4, e12559. 10.7554/eLife.12559

Spacek, M. A., Crombie, D., Bauer, Y., Born, G., Liu, X., Katzner, S., & Busse, L. (2022). Robust effects of corticothalamic feedback and behavioral state on movie responses in mouse dLGN. 11(e70469). 10.7554/eLife.70469

Speed, A., Del Rosario, J., Burgess, C. P., & Haider, B. (2019). Cortical State Fluctuations across Layers of V1 during Visual Spatial Perception. Cell Reports, 26(11), 2868–2874.e3. 10.1016/j.celrep.2019.02.045

Stüttgen, M. C., Rüter, J., & Schwarz, C. (2006). Two Psychophysical Channels of Whisker Deflection in Rats Align with Two Neuronal Classes of Primary Afferents. The Journal of Neuroscience, 26(30), 7933–7941. 10.1523/JNEUROSCI.1864-06.2006

Swadlow, H. A. (1987). Corticogeniculate neurons, cortico-tectal neurons, and suspected interneurons in visual cortex of awake rabbits: Receptive-field properties, axonal properties, and effects of EEG arousal. (57(4)), 977–1001. 10.1152/jn.1987.57.4.977

Swadlow, H. A. (1989). Efferent Neurons and Suspected Interneurons in S-1 Vibrissa Cortex of the Awake Rabbit: Receptive Fields and Axonal Properties. 62(1), 288–308. 10.1152/jn.1989.62.1.288

Swadlow, H. A., & Hicks, P. (1996). Somatosensory cortical efferent neurons of the awake rabbit: Latencies to activation via supra--and subthreshold receptive fields. 75(4). 10.1152/jn.1996.75.4.1753

Syeda, A., Zhong, L., Tung, R., Long, W., Pachitariu, M., & Stringer, C. (2022). *Facemap: A framework for modeling neural activity based on orofacial tracking* [Preprint]. Neuroscience. 10.1101/2022.11.03.515121

Temereanca, S., Brown, E. N., & Simons, D. J. (2008). Rapid Changes in Thalamic Firing Synchrony during Repetitive Whisker Stimulation. The Journal of Neuroscience, 28(44), 11153–11164. 10.1523/JNEUROSCI.1586-08.2008

Temereanca, S., & Simons, D. J. (2004). Functional Topography of Corticothalamic Feedback Enhances Thalamic Spatial Response Tuning in the Somatosensory Whisker/Barrel System. Neuron, 41(4), 639–651. 10.1016/S0896-6273(04)00046-7

Tsumoto, T., & Suda, K. (1980). Three groups of cortico-geniculate neurons and their distribution in binocular and monocular segments of cat striate cortex. Journal of Comparative Neurology, 193(1), 223–236. 10.1002/cne.901930115

Urbain, N., Salin, P. A., Libourel, P.-A., Comte, J.-C., Gentet, L. J., & Petersen, C. C. H. (2015). Whisking-Related Changes in Neuronal Firing and Membrane Potential Dynamics in the Somatosensory Thalamus of Awake Mice. Cell Reports, 13(4), 647–656. 10.1016/j.celrep.2015.09.029

Usrey, W. M., & Sherman, S. M. (2018). Corticofugal circuits: Communication lines from the cortex to the rest of the brain. Journal of Comparative Neurology, 527(3), 640–650. 10.1002/cne.24423

Usrey, W. M., & Sherman, S. M. (2019). Corticofugal circuits: Communication lines from the cortex to the rest of the brain. Journal of Comparative Neurology, 527(3), 640–650. 10.1002/cne.24423

Van Horn, S. C., Eri?ir, A., & Sherman, S. M. (2000). Relative distribution of synapses in the A-laminae of the lateral geniculate nucleus of the cat. The Journal of Comparative Neurology, 416(4), 509–520. 10.1002/(SICI)1096-9861(20000124)416:4%3C509::AID-CNE7%3E3.0.CO;2-H

Vélez-Fort, M., Bracey, E. F., Keshavarzi, S., Rousseau, C. V., Cossell, L., Lenzi, S. C., Strom, M., & Margrie, T. W. (2018). A Circuit for Integration of Head- and Visual-Motion Signals in Layer 6 of Mouse Primary Visual Cortex. Neuron, 98(1), 179–191.e6. 10.1016/j.neuron.2018.02.023

Voigts, J., Deister, C. A., & Moore, C. I. (2020). Layer 6 ensembles can selectively regulate the behavioral impact and layer-specific representation of sensory deviants. eLife, 9, e48957. 10.7554/eLife.48957

von Krosigk, M., Monckton, J. E., Reiner, P. B., & McCormick, D. A. (1999). Dynamic properties of corticothalamic excitatory postsynaptic potentials and thalamic reticular inhibitory postsynaptic potentials in thalamocortical neurons of the guinea-pig dorsal lateral geniculate nucleus. Neuroscience, 91(1), 7–20. 10.1016/S0306-4522(98)00557-0

Waiblinger, C., Borden, P. Y., & Stanley, G. B. (2022). Emerging experience-dependent dynamics in primary somatosensory cortex reflect behavioral adaptation. 13(534). 10.1038/s41467-022-28193-z

Waiblinger, C., Brugger, D., Whitmire, C. J., Stanley, G. B., & Schwarz, C. (2015). Support for the slip hypothesis from whisker-related tactile perception of rats in a noisy environment. Frontiers in Integrative Neuroscience, 9. 10.3389/fnint.2015.00053

Waiblinger, C., Reedy, A. R., & Stanley, G. B. (2026). An adaptive and flexible role for primary sensory cortex. Nature Neuroscience, 29(1), 2–12. 10.1038/s41593-025-02124-9

Waiblinger, C., Whitmire, C. J., Sederberg, A., Stanley, G. B., & Schwarz, C. (2018). Primary Tactile Thalamus Spiking Reflects Cognitive Signals. The Journal of Neuroscience, 38(21), 4870–4885. 10.1523/JNEUROSCI.2403-17.2018

Waiblinger, C., Wu, C. M., Bolus, M. F., Borden, P. Y., & Stanley, G. B. (2019). Stimulus Context and Reward Contingency Induce Behavioral Adaptation in a Rodent Tactile Detection Task. The Journal of Neuroscience, 39(6), 1088–1099. 10.1523/JNEUROSCI.2032-18.2018

Wang, Q., Webber, R. M., & Stanley, G. B. (2010). Thalamic synchrony and the adaptive gating of information flow to cortex. Nature Neuroscience, 13(12), 1534–1541. 10.1038/nn.2670

Wang, W., Andolina, I. M., Lu, Y., Jones, H. E., & Sillito, A. M. (2016). Focal Gain Control of Thalamic Visual Receptive Fields by Layer 6 Corticothalamic Feedback. Cerebral Cortex, cercor;bhw376v1. 10.1093/cercor/bhw376

Whilden, C. M., Chevée, M., An, S. Y., & Brown, S. P. (2021). The synaptic inputs and thalamic projections of two classes of layer 6 corticothalamic neurons in primary somatosensory cortex of the mouse. Journal of Comparative Neurology, 529(17), 3751–3771. 10.1002/cne.25163

Williamson, R. S., & Polley, D. B. (2019). Parallel pathways for sound processing and functional connectivity among layer 5 and 6 auditory corticofugal neurons. eLife, 8, e42974. 10.7554/eLife.42974

Wright, N. C., Borden, P. Y., Liew, Y. J., Bolus, M. F., Stoy, W. M., Forest, C. R., & Stanley, G. B. (2021). Rapid Cortical Adaptation and the Role of Thalamic Synchrony during Wakefulness. The Journal of Neuroscience: The Official Journal of the Society for Neuroscience, 41(25), 5421–5439. 10.1523/JNEUROSCI.3018-20.2021

Yang, H., Kwon, S. E., Severson, K. S., & O’Connor, D. H. (2016). Origins of choice-related activity in mouse somatosensory cortex. Nature Neuroscience, 19(1), 127–134. 10.1038/nn.4183

Ziegler, K., Folkard, R., Gonzalez, A. J., Burghardt, J., Antharvedi-Goda, S., Martin-Cortecero, J., Isaías-Camacho, E., Kaushalya, S., Tan, L. L., Kuner, T., Acuna, C., Kuner, R., Mease, R. A., & Groh, A. (2023). Primary somatosensory cortex bidirectionally modulates sensory gain and nociceptive behavior in a layer-specific manner. Nature Communications, 14(1), 2999. 10.1038/s41467-023-38798-7

